# Real-Time Axial Motion Compensation for Intravital Two-Photon Imaging of Mechanically Loaded Bone

**DOI:** 10.64898/2026.07.17.739194

**Authors:** Samantha Bratcher, Macy Mora-Antoinette, Karl J. Lewis

## Abstract

Motion artifacts present a major challenge for intravital imaging of tissues undergoing physiological movement or mechanical loading. Blurring or artificial changes in image intensity due to shifting on the z-axis reduce data reproducibility and reliability. Existing post-acquisition approaches can partially compensate for motion, but they increase experimental complexity and are not well suited for use in mechanically loaded bone. Therefore, we developed a novel method for real-time correction of axial motion during mechanical loading of bone by synchronizing the movement of the objective to the actuator. Synchronization was achieved by linking the position of the actuator piezo motor to the objective piezo motor with a user refined reduction via potentiometer. Applying axial motion correction effectively removed artificial changes in fluorescent intensity in a static fluorescent marker up to 3000με in bone as measured by similarity and average intensity before and during loading. This improvement was reflected in the improved accuracy in capture of a dynamic fluorescent calcium indicator (GCaMP6f) in osteocytes. Our system provides a user-friendly, robust framework that can be easily adapted to other mechanically loaded tissues, improving data collection and expanding the utility of two-photon imaging across a variety of biological applications.

## Introduction

*In vivo* two-photon microscopy has revolutionized the way biological tissues are studied, particularly for applications in neuroscience, developmental biology, and, more recently, bone imaging [1–3]. The ability to capture high-resolution fluorescence data deep within living tissue provides unique insights into cellular dynamics and tissue physiology. However, as with many imaging modalities, intravital two-photon imaging is highly susceptible to motion artifacts, particularly in the axial direction. These artifacts can significantly distort the data by introducing false changes in fluorescence intensity that may be erroneously interpreted as biological responses. Axial motion artifacts are especially problematic in tissues that experience deformation during imaging, such as the brain during head movements, lungs during respiration, or heart during the cardiac cycle [4–6].

Bone, despite its rigidity, is no exception to the presence of motion artifact. Mechanical loading of skeletal long bones can produce deformation on the scale of microns, introducing motion artifact that is difficult to avoid. This issue is particularly relevant in our lab, where we perform simultaneous intravital imaging and mechanical loading of the mouse third metatarsal (MT3) [1]. Our experimental setup requires precise surgical exposure and careful alignment of mechanical components, with even minor errors greatly amplifying motion artifacts. Consequently, even after rigorous training, high attrition of mouse subjects is observed due to unusable imaging data at high strain magnitudes. One approach to overcome motion artifact is to acquire images only during a rest period between each loading cycle [7]. While appropriate for long term signaling, intermittent image acquisition risks failing to capture rapid and transient calcium signals.

In studies involving simultaneous two-photon calcium imaging and moving tissues, a common solution to motion artifact is correction of collected images using post-processing. However, axial motion cannot easily be corrected in this manner because changes in fluorescence intensity due to motion are indistinguishable from activity-induced fluctuations [8]. Numerous research groups have attempted to address these challenges. One study describes an automated correction method for fast motion artifacts in two-photon imaging of awake animals [4]. Their work emphasizes the challenges of imaging dynamic tissues where displacements can introduce non-uniform distortions in the scanned images. Other research groups have experimented with two-color labeling techniques in small axial displacements, relying upon stable fluorescent markers or dyes as references [9,10]. While effective in some cases, these methods increase experimental complexity by requiring an additional fluorescent label. Furthermore, they do not actively compensate for motion, instead applying translations to image frames or correction of frame intensity from correlations with a reference volume. Another relevant example by Soulet et al. employed a system to track and correct tissue movement based on similarity to an automatically determined reference frame [11]. However, limitations surrounding image quality remain, as frames identified as distorted are simply discarded, reducing the amount of usable data. In all cases presented here, while post-processing can sufficiently improve the average quality of data from images with motion artifact, a solution that can preserve data quality and yield during continuous imaging without increasing experimental complexity is still desired.

Our work here addresses the problem of axial motion artifacts by developing a motion-correction method tailored to mechanically loaded bone imaging, building on a previously established approach from our lab that enables *in vivo* imaging of calcium dynamics in osteocytes. This motion correction device enables imaging during mechanical loading of whole bone and applies active correction[1,2]. We use a piezo-driven controller to synchronize the movements of the two-photon microscope’s objective with the mechanical loading of the bone. This synchronization ensures the imaging focal plane remains aligned with the sample, even as small deflections occur during active mechanical loading. Our system facilitates precise fluorescence measurements of calcium dynamics in bone cells, even under high strain conditions, where motion artifacts are most pronounced.

## Methods

All procedures were approved by Cornell University IACUC.

### Static fluorescent labeling with calcein

Standard female laboratory mice C57BL/6J were ordered from JAX Laboratory and aged to skeletal maturity. At 16-weeks old, mice (n=3) were injected with 200μL of calcein IP for static fluorescent labeling of osteocytes. Calcein solution was made with 0.12g calcein (Sigma; 154071-48-4) and 0.05g sodium bicarbonate in 25mL of 1x PBS. Solution was filtered for sterility under vacuum using a 0.22μm filter (MilliporeSigma; SCGP00525). Mice were imaged using two-photon microscopy 1-3 hours after injection. Osteocytes were visibly labeled with strong calcein signal.

### Dynamic fluorescent labeling with osteocyte-targeted GCaMP

Mice with osteocyte-targeted expression of GCaMP6f were generated by crossing Ai38 mice B6J.CgGt(ROSA)26Sortm95.1(CAGGCaMP6f)Hze/MwarJ (JAX Labs, strain no. 028865), which contain a Cre-dependent GCaMP6f construct under a Lox-STOP-Lox cassette [12], with DMP1-Cre mice [B6N.FVB-Tg(Dmp1-cre)1Jqfe/ BwdJ; (JAX Labs, strain no. 023047) [13]. Resulting offspring were then bred onto a C57BL/6 background. GCaMP6f acts as a calcium indicator within the cell, displaying an increase in EGFP fluorescence intensity upon binding of Ca^2+^. Previous studies with this mouse model used GCaMP3, an older variant, and validated fluorescent expression in cortical osteocytes [14]. Our substitution with GCaMP6f was motivated by its design as an ultrasensitive protein calcium sensor with faster kinetics and greater dynamic range than alternative GCaMPs [12].

### In vivo loading of the third metatarsal

Mouse MT3s were surgically exposed and placed into a custom made stainless steel *in vivo* three-point bending device as previously described [1,2]. Briefly, mice were initially placed under anesthesia in a small container with 1L/min flow rate and 3% isoflurane. Mice were then transferred onto the loading device stage with a heating pad and nose cone and the isoflurane reduced to 2%. Tendons overlying the MT3 were surgically removed, while adjacent blood vessels were kept intact. The fulcrum pin was inserted between the MT3 and underlying tissue and the bracket attached overlying the pin. The paw was placed in a room temperature DPBS bath for the duration of the experiment. Cyclic haversine position-controlled loading was applied at 1Hz for 60 seconds to previously calibrated peak strains of 1000-, 2000- and 3000με. These levels encompass the range of strains that have been reported during physiological activities from *in vivo* strain gauge studies, with strains up to 2000με characteristic of habitual activities, while strains on the order of 3000με are seen in extreme activities [15,16].

### Correction of motion artifacts at high loading displacements

Axial motion correction was achieved by moving the microscope objective synchronously with the position-controlled loading regime in a custom circuit configuration (Figure 1). The analog signal of the loading actuator controller (Physik Instrumente, E-661) was sent to the objective controller (Thorlabs, PPC001). However, the objective only needs to move a small fraction of the actuator distance on par with the small deflections seen in the bone. Therefore, a potentiometer (20kΩ; Bourns-3590P-2-203L) was added to allow the user to manually increase the resistance and reduce the signal amplitude as needed. A second potentiometer (2kΩ; Bourns-3362P-1-202LF) was added as a second constant input to ensure the signal will maintain the same average value as the user increases the resistance of the first. Buffered operational amplifiers (op-amps, elements that contain their own power sources) were included to prevent signal attenuation that may occur throughout the circuit. Grounded capacitors (1μF) at the two inputs were used for fine-tuning by removing high-frequency and low-level noise that might exist in the system. An op-amp inverter (10 kΩ) was added, as the piezo crystal of the loading actuator and the objective are oriented in reverse. The op-amps were consolidated using 2 dual-circuit op-amps (Texas Instruments, TLV272 Rail-to-Rail). The final two resistors (10 kΩ) provide safety factors should the system ever be short circuited.

**Figure 1.**
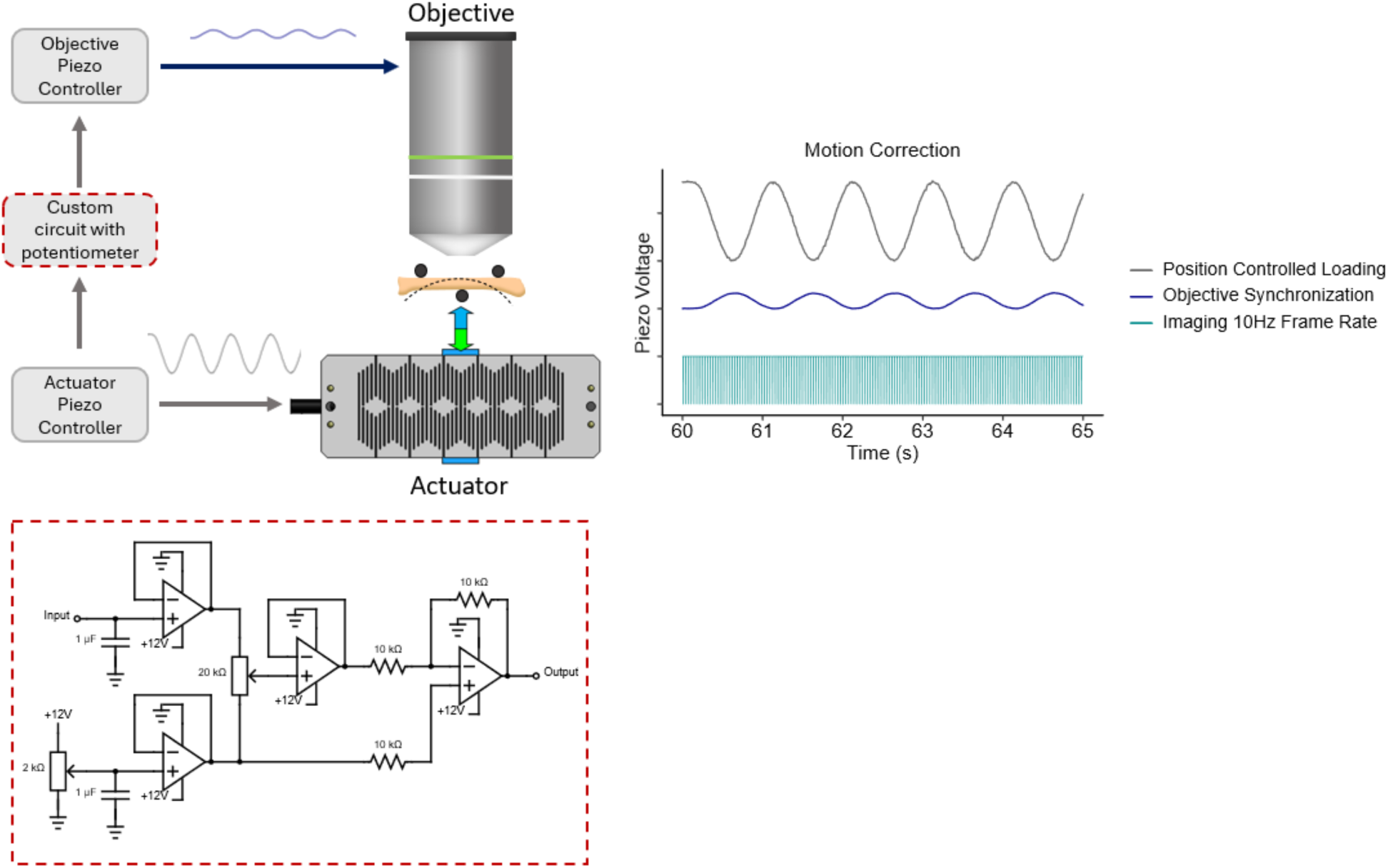
A diagram showing how a second piezoelectric controller can move the objective synchronously with the loading actuator, synchronizing the objective movement with that of the small deflections on the bone during compressive loading. (Left top) The actuator amplitude is reduced with a user-defined variable resistance using a potentiometer. (Left bottom) The circuit diagram connecting the two controllers for both the loading actuator and the objective. The input is the signal to directly control the loading actuator, and the output is the user defined reduction of the signal for the objective. (Right) Data collected from the ThorSync software showing the piezo voltages of the different signals, including the input (loading device), output (objective), and frame acquisition.

### Intravital imaging with two-photon microscopy

*In vivo* static and dynamic fluorescence signals were visualized with a two-photon microscope (Thor Labs Bergamo II; Newton, NJ) and a 20x water immersion objective (XLUMPLFLN, Olympus). Osteocytes were imaged at the dorsal mid-shaft region of the MT3 directly above the fulcrum pin beneath the bone and beneath the periosteal surface, verified with visible autofluorescence from the bone surface. The ROI was 512 x 512 pixels with 2x zoom leading to a pixel size of 0.545 x 0.545μm. Excitation was at 920nm wavelength from a tunable Ti- Saphire laser (Coherent, Chameleon Discovery NX). For calcein labeling, the power selected was such that the final output from the objective was 10mW, and a PMT with a 490-560nm bandpass filter and a voltage gain of 0.6V was used for detection. For GCaMP6f labeling, the power was between 30-35mW and a PMT voltage gain of 0.7V. Time series images were acquired by averaging 3 frames with a sampling rate of 29.4Hz, leading to a final frame rate of 9.8fps. Across an approximate 150 second period, 1500 8-bit TIF images were collected, with 60 seconds non-loading, 60 seconds loading, and 30 seconds resting.

### Error analysis with structural similarity index

The Structural Similarity Index Measure (SSIM) is a method for measuring the similarity between two images. Unlike traditional measures like Mean Squared Error (MSE) or Peak Signal-to-Noise Ratio (PSNR), which consider absolute errors, SSIM is designed to model the human visual system’s perception of image quality using a weighted combination of the changes in luminance, contrast, and structure. The SSIM index between two images x and y is defined according to equation 1.

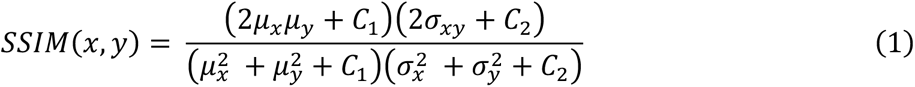

Where μ_x_ and μ_y_ are the means of the images x and y, σ_x_^2^ and σ_y_^2^ are the variances of the images x and y, σ_xy_ is the covariance of the images x and y. C_1_ and C_2_ are constants to avoid instability when the means and variances are very small. SSIM equals 1 when two images are identical (perfect similarity) and equals 0 when they share no similarity. Since calcein is a static fluorescent marker, it can be assumed that changes in fluorescence during mechanical loading that cause reductions in image similarity are likely due to motion artifacts. Comparing images across a loading cycle to images under a non-loaded reference could therefore be used to determine the prominence of the motion artifact in the system. SSIM was computed in MATLAB (MathWorks).

### Fluorescent intensity analysis

Intensity measurements were performed by postprocessing image time series using ImageJ (1.54f NIH, Java 1.8.0_322). In-plane movements in the x and y were removed using a template matching plugin. ROIs were selected with a semi-automated process. Image series were projected across the z with mean intensity, contrast enhanced, and smoothed by a gaussian blur with sigma of 1. Cell ROIs were generated by user defined thresholding and automatic particle analysis filtering ROIs within 150-650 pixels^2^. Final adjustments were made by manually addition of ROIs that were not properly captured by thresholding or deletion of ROIs of cells which appear to shift out-of-plane during loading. An ROI indicating the background autofluorescence was chosen in an area where no cells were present. The average intensity of each ROI was collected across all 1500 time series frames and output as a csv.

In MATLAB, ROIs with average intensities lower than the mean background intensity plus 3 standard deviations were excluded as background osteocytes. The intensities for each cell of interest were then detrended linearly to correct for photobleaching and normalized to the mean intensity for that cell over a 40 second period prior to loading. Peaks and troughs were identified as having a prominence of 0.01. Fluorescent intensity amplitudes across the non-loaded and loaded intervals were computed by subtracting coupled troughs from peaks. Normalization of fluorescent intensity was computed as *F_load_*/*F_non_*_–*load*_ with static values *F_non_*_–*load*_/*F_non_*_–*load*_ = 1 being used as reference controls. F_load_ indicates the amplitude during loading and F_non–load_ indicates the amplitude without loading. ROIs in time series from GCaMP6f mice were additionally classified as responsive if fluorescence intensity increased by more than 25% during loading compared to non-loading conditions.

### Statistical analysis

The effect of the corrected versus uncorrected motion on the SSIM of a static fluorescent marker was determined using a paired two-tailed T-test with alternative hypotheses of uncorrected data having lower SSIM. Change in fluorescent intensity during loading of a static fluorescent marker was determined using a paired two-tailed T-test with alternative hypotheses that fluorescence during loading is higher than the initial static portion. Statistical tests were performed in Prism (GraphPad, Version 11).

## Results

### Calibrations for male and female GCaMP6f mice

To determine the actuator displacements required to reach physiological strain values spanning the range of 1000-3000με, calibrations using three-point bending were performed on previously dissected GCaMP6f mouse paws. Strain and load maintain a consistent linear relationship with displacement within male and female groups (Figure S1). The displacement required to reach a strain value of 3000 με is near 40μm for male and female groups with expected average maximum loads to be greater in male mice than female mice (Figure S1). These values are slightly higher than have been previously reported for GCaMP3 mice [1]. As a result, loading at 2000με and 3000με produced noticeable motion artifacts, motivating the development of a motion artifact correction system that was not required in prior experiments.

### Successful removal of motion artifacts allows for higher loading displacements

In order to determine if motion artifact is successfully removed while loading at higher displacements, we used a static fluorescent marker, calcein, for reference. As a static marker, changes in the fluorescent intensity during loading are likely due to axial motion artifact leading to artificial fluorescent changes. Mice were injected with calcein intraperitoneally and imaged 1-3 hours later with metatarsal loading at 1000με (Supplemental Video 1), 2000με (Supplemental Video 2) and 3000με (Supplemental Video 3). Images were captured both with and without motion correction.

The synchronization of the objective with the loading regime successfully removed the motion artifact seen at higher load displacements (Figure 2). Without motion correction, the ROIs tend to change over the duration of a single loading cycle, and motion artifact in the z-plane worsens with increasing applied strain. However, with the objective synchronized using the user-defined signal reduction of the position-controlled loading regime, little to no changes are visible in the ROI as indicated by similarity scores close to 1. This is further highlighted in the SSIM during the 60s loading period (Figure 3). As maximum displacement was reached, the SSIM reduced, with larger reductions in SSIM aligning with higher applied load. Once motion correction is applied, significant changes from the reference frame are removed as observed by the average SSIM value remaining closer to 1.

**Figure 2.**
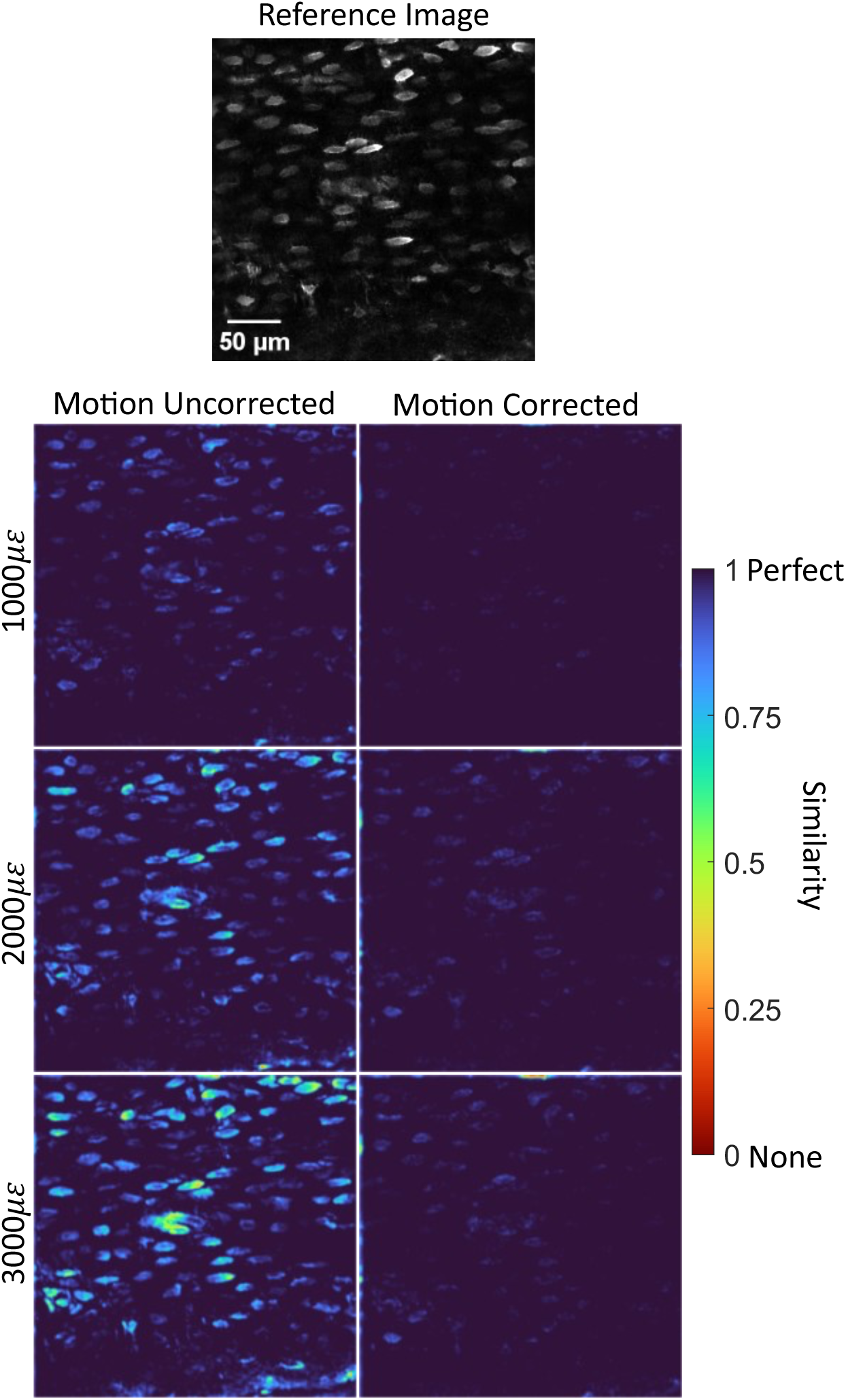
A heatmap of SSIM (similarity between loaded and non-loaded images) in calcein injected mice. Uncorrected motion is shown on the left and corrected motion is shown on the right. Areas with high differences (i.e. low similarity) are visible at 2000με and 3000με, exemplifying how intensity changes are falsely created when motion is not corrected while correcting motion removes these artificial variations.

**Figure 3.**
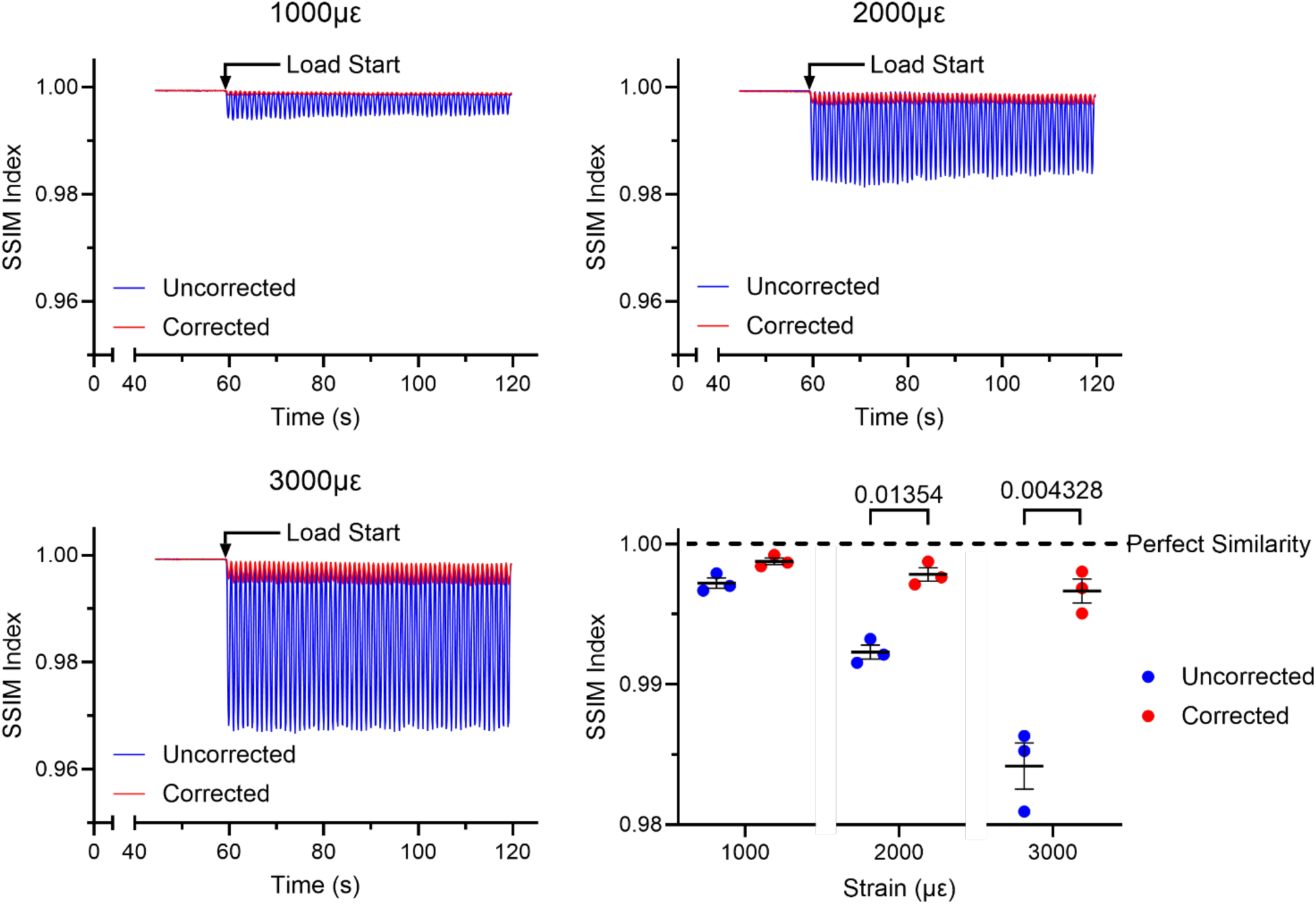
Average SSIM (Similarity index) during the 60 second loading period in calcein injected mice. Change from the static reference frame followed the cyclic pattern of loading and increased with applied strain. With motion correction from objective synchronization, all significant changes to the focal plane are removed, increasing the average similarity to near 1 during mechanical loading. n=3. Error=SEM.

We additionally quantified the artificial changes in fluorescence that may be produced if axial motion is not properly corrected by comparing the fluorescent intensity during loading to the non-loading portion. At all strain levels, the average uncorrected fluorescent intensity of all cells during loading is greater than the non-loaded (Figure 4). However, applying motion correction removes changes in fluorescent intensity during mechanical loading. Normalizing the uncorrected ΔF/F by the corrected ΔF/F further reveals drastic artificial increases in uncorrected image data. Thus, motion artifact correction using objective synchronization will allow for successful capture of calcium dynamics without false positive results that in-and-out of plane movement at high loading strain may introduce.

**Figure 4.**
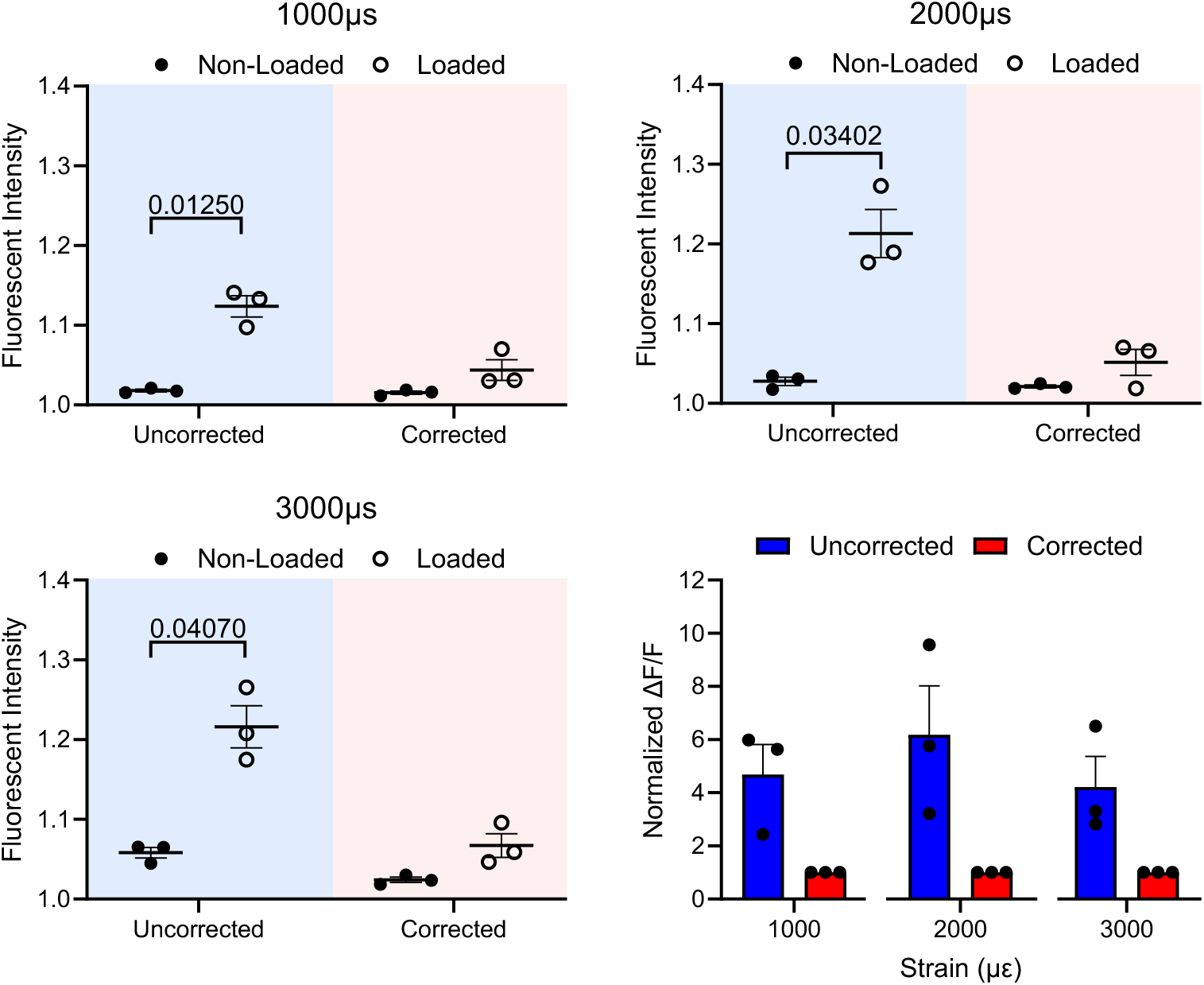
The average fluorescent intensity of all cells in calcein injected mice show an artificial increase in intensity during loading at 1000με, 2000με, and 3000με. Correcting for motion removes changes in the static calcein signal, creating a stable image. When normalizing the uncorrected ΔF/F by the corrected ΔF/F, we saw drastic artificial increases in fluorescence in comparison when motion is not corrected. n=3. Error=SEM.

### Robust calcium signals within osteocytes are collected with GCaMP6f mice

By eliminating motion artifact, calcium signals may be more accurately captured at high mechanical strains. We validated this by performing MT3 loading on a 16-week-old female mouse expressing GCaMP6f in which the fluorescent intensity within osteocytes varied dynamically with intracellular calcium concentration in response to strain magnitude. Motion correction enabled us to capture prominent changes in fluorescent intensity (Figure 5). In motion-uncorrected time series, cell ROIs exhibited sharp, uniform decreases in fluorescent intensity that mirror increases in actuator displacement. In contrast, cell ROIs in motion-corrected time series fluctuate more smoothly about their normalized baseline intensities with varying peak amplitude indicative of physiological calcium signaling. Motion correction also substantially increased the number of analyzable cells. Only 8 osteocytes were considered viable in the uncorrected time series, whereas 53 usable cells were identified following correction. Among these, 100% of viable uncorrected ROIs were classified as responsive, compared with 98% responsiveness in corrected time series. Collectively, these changes may be accurately attributed to osteocyte response to mechanical loading rather than motion artifact due to the effective removal of shifting in the z axis by motion correction.

**Figure 5.**
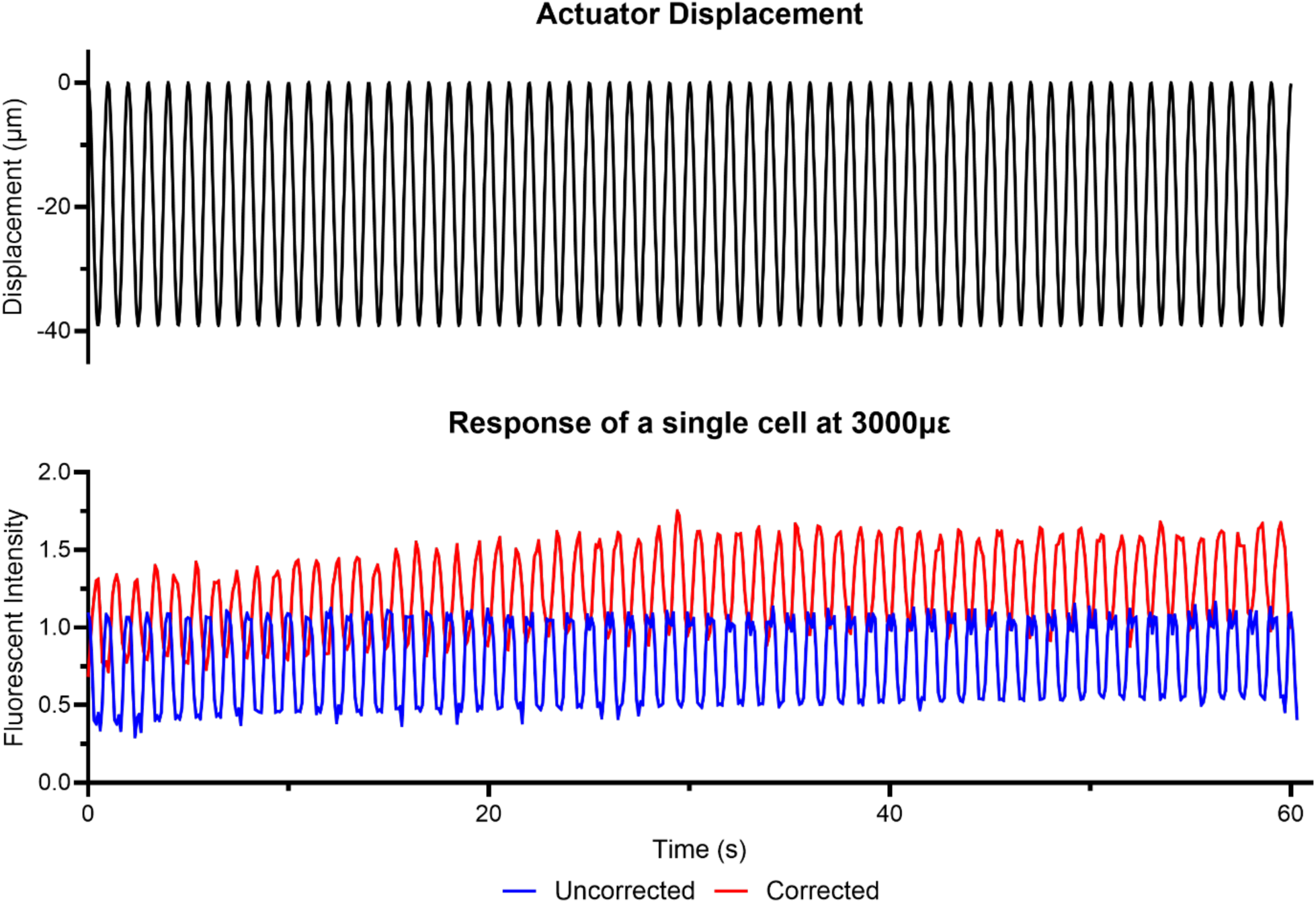
Comparison of fluorescent intensity of a representative single cell in a female GCaMP6f mouse. Fluorescent signal in an uncorrected image series decreases dramatically alongside actuator displacement. In contrast, a smoother calcium signal in response to load is captured with motion correction, varying about a normalized baseline intensity.

## Discussion

Axial motion on the scale of microns is difficult to avoid during intravital imaging of mechanically loaded bone. Resulting artifacts reduce accuracy and reliability of collected data. To address this, we developed a system that uses two independent controllers to operate separate piezo devices. The signal driving the piezo actuator, which applies mechanical load to the bone, is sent to the piezo controller of the microscope objective with a user defined reduction in amplitude. By synchronizing the analog signals between these controllers, we ensured the objective follows the small displacements of the bone, effectively removing artificial changes in fluorescent intensity caused by shifting of the focal plane. This approach enabled highly accurate recordings of osteocyte response to mechanical stimulation, even at high strain levels.

As a mechanoresponsive cell, it is crucial osteocytes are studied both under mechanical load and *in vivo* to fully understand their physiological behavior. While osteocytes produce autonomous calcium signals under static conditions, they respond actively to mechanical stimulation in a strain-dependent manner [1,7,17]. Consequent conversion of tissue strain into biochemical signals through mechanotransduction is highly dependent on the immediate microenvironment. Matrix-binding proteins (e.g., proteoglycans, integrins) form functional attachments between osteocyte cell membranes and the matrix wall that are critical for fluid activation of the cell [18–20]. In addition, the morphology of the lacunae and canaliculi housing the cell body and dendritic processes plays a critical role in tissue strain and interstitial fluid flow [21–23]. Finally, osteocyte mechanosensitivity is further mediated by a multitude of channels and receptors involved in Ca^2+^ signaling dynamics and downstream pathways [24–27].

Although osteocyte response to mechanical loading has been previously accomplished, here we present a critical improvement for axial motion artifact and distinctly improve data collection efficiency. This is especially beneficial in experiments where differences in bone mechanical properties associated with age, disease, or mouse model may require larger tissue displacements for calibrated strains[1]. For example, we observed that mice expressing GCaMP6f required greater displacement during mechanical loading of the MT3 compared to those expressing GCaMP3. These larger displacements introduced more pronounced motion artifacts than previously observed, which were further exacerbated by the sensitivity of the experimental setup. Achieving a consistent and stable surgical setup of the MT3 can be challenging even after thorough training. Variations in bone diameter and the presence of soft tissue between the bone and isolating pin can reduce stability. In parallel, machined components of the loading apparatus require precise alignment and operate within tight tolerances. Within our lab, high-strain loading experiments without motion correction have resulted in as low as 20-30% successful data. Incorporating motion correction improves the quality and reproducibility of collected data and reduces the total number of experimental animals needed.

The synchronization system presented here provides a robust and proactive solution to axial motion by enabling real-time correction and prevention of motion artifacts during image acquisition. Axial motion is more difficult to correct than lateral motion due to changes in image intensity, focus, and feature appearance, requiring correlation to a three-dimensional reference volume. As a result, conventional post-processing approaches can only reduce data uncertainty by computationally adjusting the frame intensity or rejecting frames with too much motion. These attempts to correct artifacts can introduce other inaccuracies through interpolation of signal or error in registration of features. In contrast, the system developed here rectifies axial motion at its source to preserve the integrity of raw data. Furthermore, it increases data yield without additional computational time and power by negating the need for correction in post-processing.

Active motion correction during simultaneous imaging and loading represents a significant advancement in the field of bone imaging, particularly for studies involving mechanical loading and calcium signaling. By minimizing motion artifacts during *in vivo* imaging, we can reduce animal usage and more accurately assess the biological response of bone cells to high levels of mechanical stress, offering new insights into bone physiology and mechanotransduction. Additionally, our system provides a user-friendly, robust framework that can be easily adapted to other mechanically loaded tissues, expanding the utility of two-photon imaging across a variety of biological applications.

## Acknowledgements

We thank Karly Hooper for maintaining mouse colonies for use in these experiments, along with the entire CARE staff in Weill Hall. We also thank Daniel Rivera for assistance with the design and troubleshooting of the motion correction circuitry. This work was supported by the Department of Biomedical Engineering at Cornell University, the Mong Family Foundation at Cornell University, and the Ford Foundation by the National Academies of Sciences, Medicine, and Engineering, NIH S10OD023466.

## Supplemental Methods

### Calibration of applied strain to the mouse third metatarsal

Before loading, calibrations were conducted with digital image correlation (DIC) to determine position-controlled loading regimes with corresponding strain values. Hind paws from 3-6 male and female mice were amputated and stored in saline soaked gauze at -20°C until calibration. Paws were then thawed at room temperature in DPBS. The MT3 was surgically isolated as described elsewhere [2] and a 100μm diameter cylindrical fulcrum pin was inserted just underneath, separating the bone from surrounding muscle fascia and soft tissue. A bracket was then positioned over the foot and connected with a screw to the load cell and actuator below. A stepladder loading regime under reverse 3-point bending compression taking 5μm incremental steps with 2 second intervals holding load constant until reaching 60μm displacement. Once maximum displacement was reached, load was released in the reverse step pattern. For DIC, 26μm diameter Microspheres (Polysciences, 18241-2) were applied to the MT3 cortex. Images were taken during loading with a 10x objective at 30 fps with 410nm epifluorescence and optics from a two-photon microscope (Thor Labs, Bergamo II). Changes in displacements of the beads were computed in MATLAB (MathWorks) to determine strain values on the surface of the bone.

## Supplemental Figures

**Figure S1.**
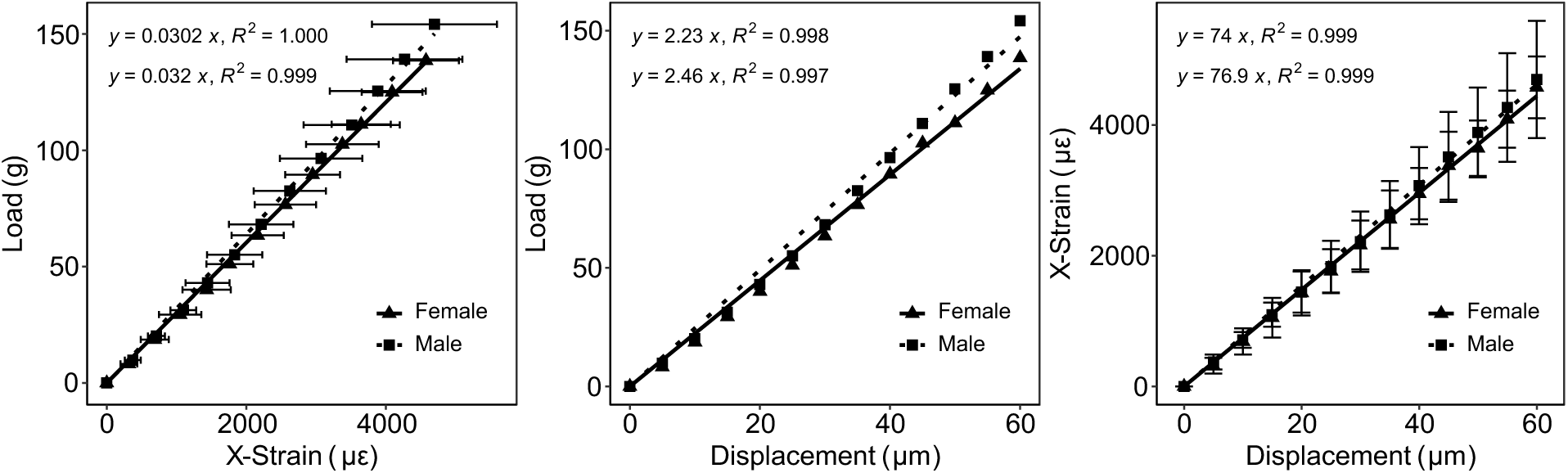
Resulting calibrations between displacement, load, and strain in the MT3 of female and male GCaMP6f mice. For calibrations, linear regressions were conducted using RStudio (Posit, v.3.6) with a controlled intercept through the origin. Regressions were made with X- strain vs displacement as well as X-strain vs load to compute applied displacement values for position-controlled loading and the expected average max load.

**Video S1.**
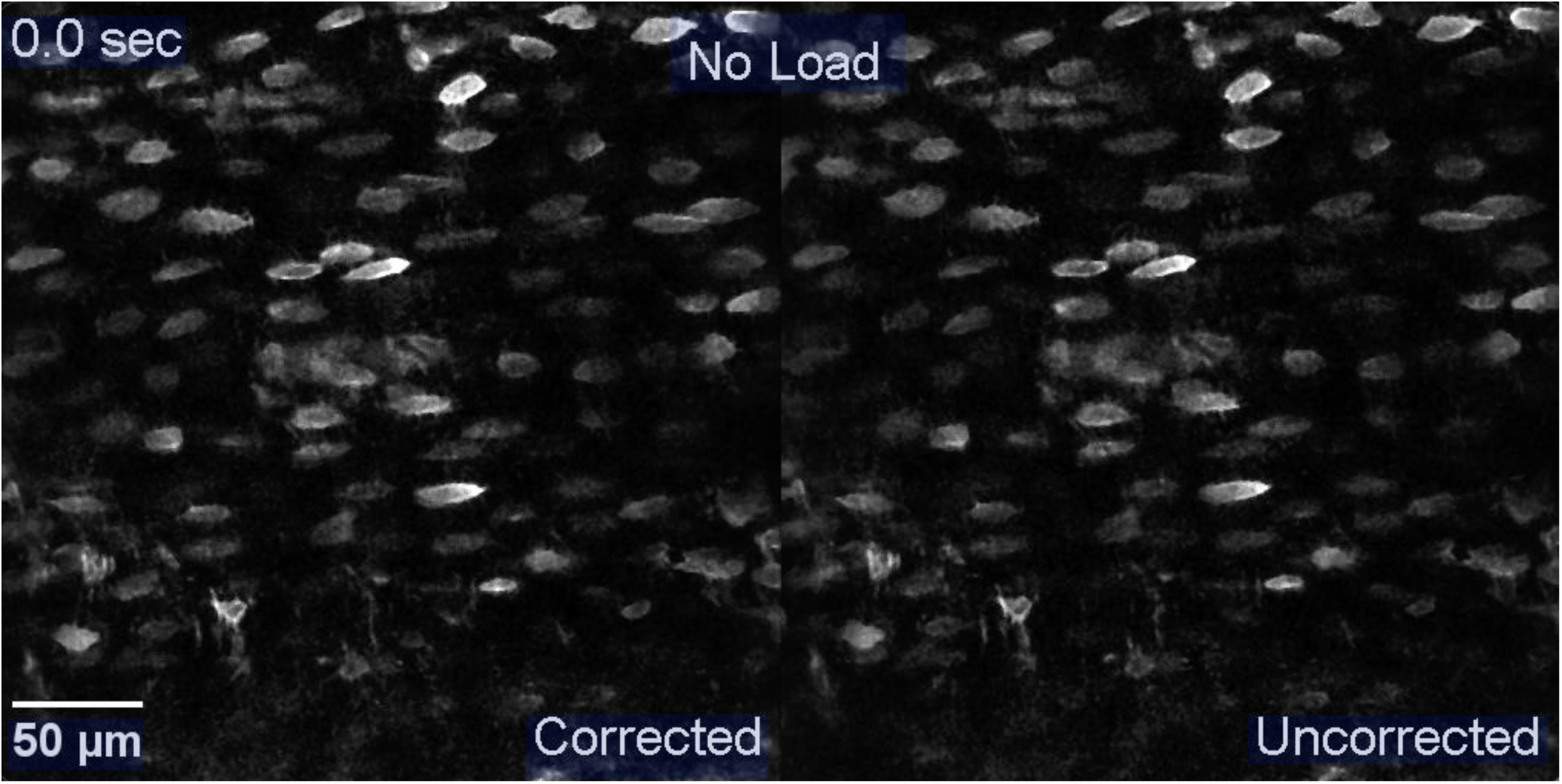
Timelapse imaging of osteocytes labelled with calcein during mechanical loading at 1000με without (right) and with (left) motion correction. Upon initiation of loading, some motion in the z-axis is noted in the motion-uncorrected time series, as evidenced by fluctuations in fluorescent signal intensity. In contrast, the motion-corrected time series remains stable.

**Video S2.**
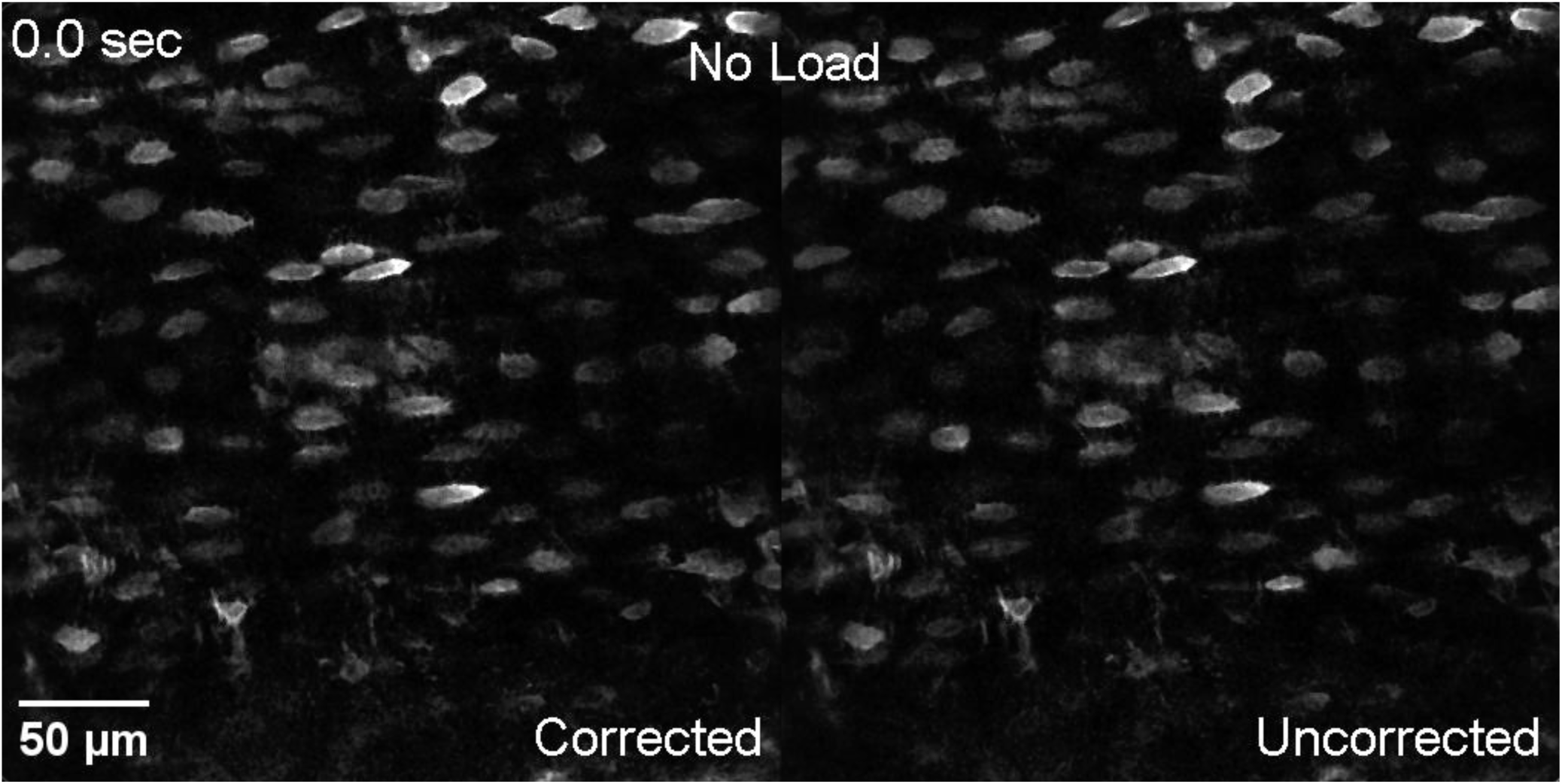
Timelapse imaging of osteocytes labelled with calcein during mechanical loading at 2000με without (right) and with (left) motion correction. Upon initiation of loading, motion in the z-axis is observed in the motion-uncorrected time series, as evidenced by drastic changes in fluorescent intensity. In contrast, the motion-corrected time series remains stable.

**Video S3.**
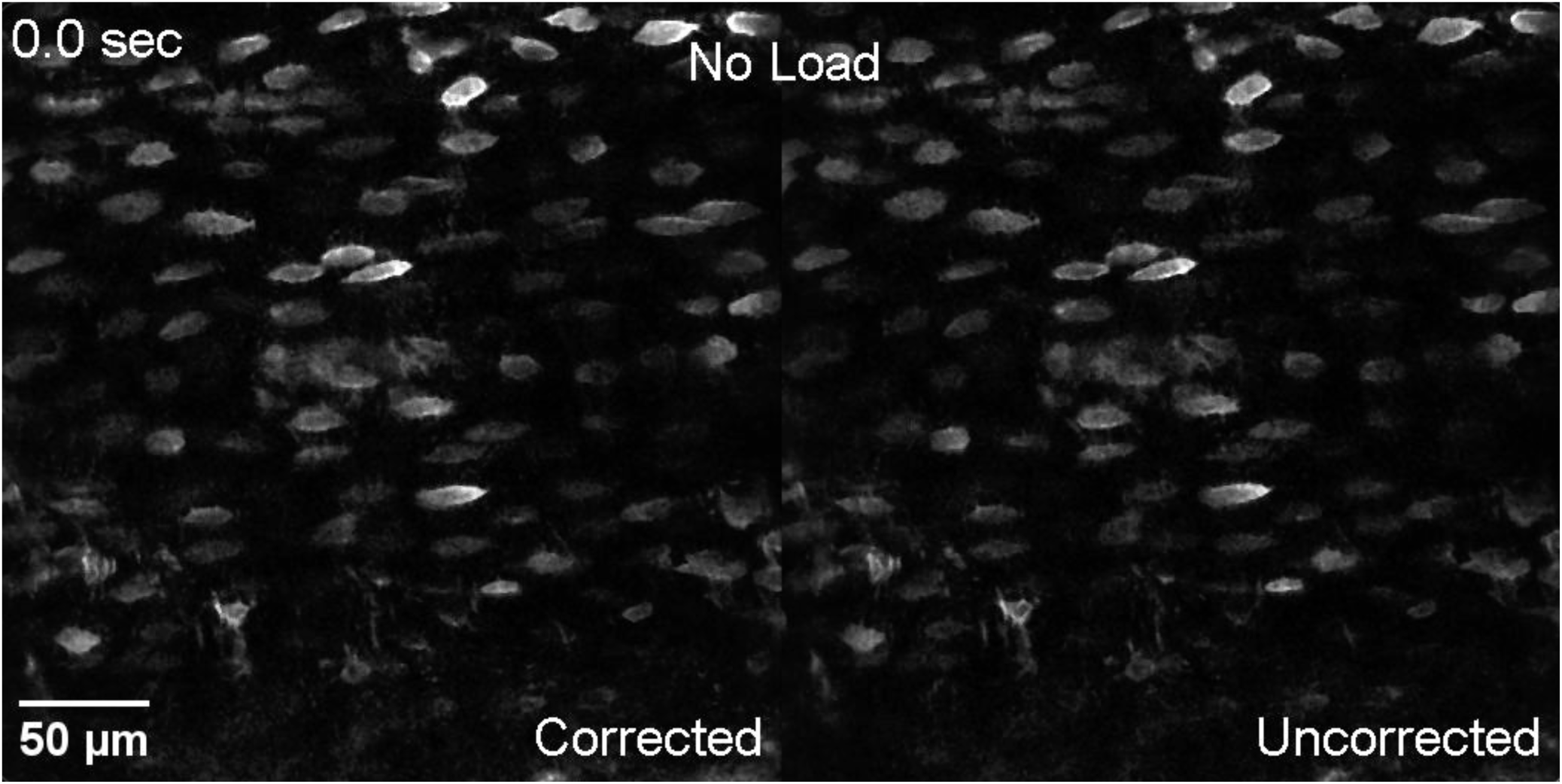
Timelapse imaging of osteocytes labelled with calcein during mechanical loading at 3000με without (right) and with (left) motion correction. Upon initiation of loading, a large degree of motion in the z-axis is apparent in the motion-uncorrected time series, with many cells shifting entirely in-and-out of the imaging plane. In contrast, the motion-corrected time series remains stable, with only minor changes in intensity of some cells.

